# Notch signalling mediates secondary senescence

**DOI:** 10.1101/554741

**Authors:** Yee Voan Teo, Nattaphong Rattanavirotkul, Angela Salzano, Andrea Quintanilla, Nuria Tarrats, Christos Kiourtis, Miryam Mueller, Anthony R. Green, Peter D. Adams, Juan-Carlos Acosta, Thomas G. Bird, Kristina Kirschner, Nicola Neretti, Tamir Chandra

## Abstract

Oncogene induced senescence (OIS) is a tumour suppressive response to oncogene activation that can be transmitted to neighbouring cells through secreted factors of the senescence associated secretory phenotype (SASP). Using single-cell transcriptomics we observed two distinct endpoints, a primary marked by Ras and a secondary by Notch. We find that secondary senescence *in vitro* and *in vivo* requires Notch, rather than SASP alone as previously thought. Currently, primary and secondary senescent cells are not thought of as functionally distinct endpoints. A blunted SASP response and the induction of fibrillar collagens in secondary senescence compared to OIS point towards a functional diversification.

**One Sentence Summary:** Notch signalling is an essential driver of secondary senescence with primary and secondary senescence being distinct molecular endpoints.

## Main Text

Cellular senescence is a stress response, resulting in a stable cell cycle arrest, and has been implicated in tumour suppression, ageing and wound healing (*1*–*4*). Aberrant activation of the Ras oncogene triggers oncogene-induced senescence (OIS), conferring a precancerous state (*5,6*). OIS is an *in vivo* tumour suppressor mechanism preventing transformation and halting tumour growth (*7,8*) with the p53 and Rb/p16 pathways acting as major mediators of senescence induction and maintenance (*6,9*). OIS is characterised by multiple changes in phenotype, such as heterochromatic foci (*9*–*13*) and the senescence-associated secretory phenotype (SASP, *14*–*16*). Through the secretion of extracellular matrix proteases and interleukins and chemokines, OIS cells recruit immune cells, mediating their own clearance. However, SASP has been implicated in cancer initiation (*17*) by creating an inflammatory pro-tumorigenic microenvironment. Moreover, SASP factors play a role in cellular reprogramming (*18,19*) and contribute to ageing and tissue degeneration (*20*–*21*). SASP acts in a paracrine fashion to induce secondary senescence in surrounding cells and mediates tissue re-organisation. Secondary senescence occurs in cells not directly receiving the stress insult^22^ with paracrine secondary senescence being mediated by secreted SASP factors. Paracrine secondary senescence is thought to enhance immune surveillance and to act as a fail-safe mechanism minimising chances of retaining damaged cells (*16,22,23*). More recently, ectopic Notch pathway activation has been implicated as an intermediate, unstable phenomenon during primary senescence induction, resulting in a distinct secretome. However, Notch as an endpoint for primary senescence and in secondary senescence mediation as a complementary pathway to paracrine secondary senescence remains undescribed.

Here we use single cell transcriptomics to decipher the heterogeneity within OIS populations. Our single cell experiments reveal two distinct transcriptional end-points in primary senescence, separated by their activation of Notch, resembling but distinct from ectopic Notch induced senescence (NIS). We find secondary senescent cells to uniformly progress to an end-point characterised by Notch activation *in vivo* and *in vit*ro. Finally, we confirm Notch mediated senescence as an essential mediator of secondary, juxtacrine senescence in OIS. Using the power of single cell transcriptomics we deconvolute cell fate decisions after oncogene activation which might confer differences in transformation and rejuvenation potential.

## Primary and secondary senescence have distinct transcriptomes

To investigate the dynamic changes, trajectory and cell-cell heterogeneity in OIS, we performed a single-cell RNA-seq time course before and after 2, 4 and 7 days of RasV12 induction using H-RasG12V-induced IMR90 (ER:IMR90) senescent fibroblasts as previously described (*24*) and Smart-Seq2 protocol (*25*) (Fig.1a). After stringent filtering (Fig.1b, Supplementary Fig.1a-d, Supplementary Table 1), we obtained a final cell count of 100/288 for day 0, 41/96 for day 2, 42/96 for day 4 and 41/288 for day 7 for downstream analysis (Supplementary Fig.1d). To confirm a senescence phenotype at day 7, we profiled Bromodeoxyuridine (BrdU) incorporation (37/390 cells (9%)), SAHF (265/390 cells (68%)) and senescence associated beta-galactosidase (SA-Beta Gal) (428/523 cells (82%)) (Supplementary Fig.1e). To assess time-dependent changes in the transcriptome, we ordered cells along a pseudo-temporal trajectory based on differential gene expression between growing and senescence (adjusted p<0.05, Table 1), by single cell differential expression (SCDE, Fig.1c) (*26*). Using Monocle2 (*27*) we found a continuous progression from growing to senescence, with days 2 and 4 cells as intermediates. The Monocle2 trajectory revealed two distinct senescent populations (Fig.1c), suggesting two facultative, alternative endpoints. To determine whether RasV12 activation led to the split into two senescence populations (Fig.1c), we overlaid RasV12 expression onto the monocle plot (Fig.1b,c, Supplementary Fig.1f,g and Supplementary Table2). RasV12 expressing cells (Fig.1c, Ras+, round symbols) progressed to both senescence endpoints with a 21:4 skew towards the cluster designated OIS. In contrast, fibroblasts without detectable RasV12 expression uniformly progressed to the cluster tentatively designated secondary senescence, suggesting it as the obligate endpoint (cross symbols =Ras-, Fig. 1c, Fisher’s exact test=1.64×10^−6^). Our inability to detect RasV12 in a subset of senescent cells suggests that senescence was induced as a secondary event. We verified HRAS as one of the top predicted upstream regulators for the senescence top, but not bottom, population (p=3.1×10^−34^) using Ingenuity pathway analysis (IPA, Qiagen, Supplementary Fig.1h). We confirmed a senescence phenotype for both populations by upregulation of key senescence genes (Fig.1d) cyclin dependent kinase 1a (CDKN1A), cyclin dependent kinase inhibitor 2b (CDKN2B) and SASP factors interleukin 8 (IL8), interleukin 6 (IL6) and interleukin 1B (IL1B (p<0.05 for all genes, Fig.1d). To verify two major senescence populations transcriptome-wide, we used a consensus clustering approach, SC3 (*28*), with the number of clusters determined by silhouette plot(*29*) (Supplementary Fig.1i). SC3 detected two senescence clusters which largely overlapped with the subpopulations obtained by Monocle2 (Cluster 1 16/21 or 76% RasV12+ cells, Cluster 4 11/15 or 73% RasV12- cells), supporting the notion that the split into two senescence populations is based on the absence or presence of RasV12 (Fig.1e). To verify that populations observed are due to primary OIS and secondary senescence, we co-cultured ER:IMR90 with IMR90:GFP fibroblasts (1:10), where secondary senescence is induced in IMR90:GFP positive cells (*22*). We generated single-cell RNA-seq transcriptomes before and 7 days after RasV12 activation, using the 10x Genomics Chromium (Fig.1f).

**Fig. 1.**
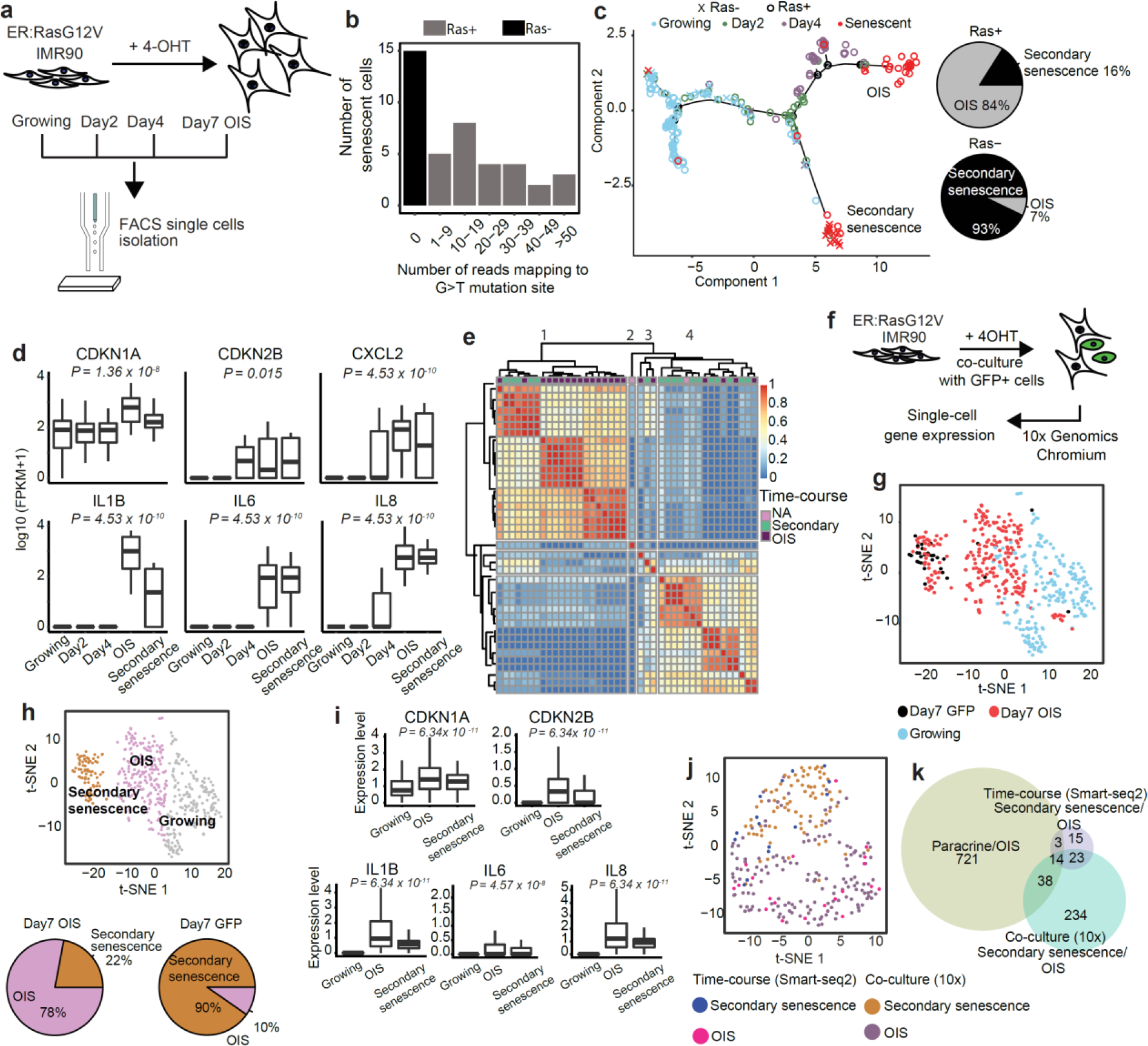
Secondary senescent cells only partially resemble paracrine induced senescence. **a.** Schematic representation of the time-course experiment. **b.** Number of senescent cells with reads mapping to the G>T mutation site of *Ras* gene. **c.** Monocle2 was used to order single cells from four time points based on the 680 DE genes (p-values < 0.05) between growing and senescent cells. Cells were annotated with the presence of the mutated *Ras* gene and the pie charts showed the percentage of Ras+/Ras- cells in the top and bottom clusters. **d.** Box plots for the expression of senescence-associated genes in the time-course experiment. The top and bottom bounds of the boxplot correspond to the 75 and 25th percentile, respectively. Differential expression was assessed using SCDE. **e.** Unsupervised clustering using SC3 cluster senescent cells. Cells were annotated as either OIS (top senescence branch, purple), secondary senescence (bottom branch, green) or NA (neither in the bottom nor top branch, pink). **f.** Schematic representation of the co-culture experiment. **g.** tSNE visualization of 480 single cells (240 Day7 OIS cells, 33 Day7 GFP cells and 207 growing cells). **h.** tSNE visualization of single cells grouped into 3 clusters. **i.** Box plots for the expression of senescence-associated genes in the co-culture experiment. The top and bottom bounds of the boxplot correspond to the 75 and 25th percentile, respectively. Differential expression was assessed using SCDE. **j.** Integration analysis of the two senescence clusters from both time-course and co-culture experiments. **k.** Overlap of DE genes between paracrine/OIS, time-course and co-culture experiments DE genes.

Senescence was confirmed on sorted populations by qPCR (Supplementary Fig.1j) and SA-beta gal staining for primary and secondary senescent cells (Supplementary Fig.1k). Cells were annotated based on GFP, RasV12 expression and the G>T mutation of *Ras* gene (Fig.1g). We identified three distinct clusters using Seurat, namely growing (blue dots), secondary senescence (GFP positive, black dots) and OIS (RasV12 positive, red dots), with significant enrichment for the OIS and secondary senescence populations (Chi-squared test, p=4.1×10^−14^, Fig.1h). The secondary senescence cluster also contained a detectable, minor population of RasV12 expressing cells. This mirrors our earlier findings, confirming two facultative senescence endpoints for primary RasV12 senescent cells with GFP positive secondary senescent cells showing a more uniform distribution. Senescence genes were upregulated in both senescent clusters, including CDKN1A, CDKN2B and IL8 (Fig.1i, Table1) and long-term stable cell cycle arrest confirmed at 21 days post co-culture (Supplementary Fig.1l). When overlaying transcriptomes of the time course and the co-culture experiments, a significant number of cells identified as OIS and secondary senescence (GFP and part of RasV12) clustered together (Fig.1j, Chi-squared test, p<0.05). Notably, the co-clustering by senescent signatures was achieved despite the data being generated on two different platforms, 10x and Smart-seq2. In summary, we identified two major transcriptional endpoints in primary OIS, whereas secondary senescent cells were uniformly assigned.

Paracrine senescence is thought to be the main effector mechanism for secondary or cell extrinsic senescence induction (*16,22*). To test if the secondary senescence is explained by a paracrine signature, we overlaid publicly available bulk RNA-Seq data (*22*). While we found a significant overlap with a paracrine signature (Hypergeometric test: Paracrine/OIS and time-course secondary senescence/OIS (Ras-/Ras+) p<0.001; Paracrine/OIS and 10x secondary senescence/OIS p<0.001, 10x secondary senescence/OIS and time-course secondary senescence/OIS (Ras-/Ras+) p<0.001, Fig.1k, Supplementary Table 3), a large fraction of genes shared between our two single cell experiments remained unexplained, suggesting the involvement of additional pathways in secondary senescence.

### The transcriptome of secondary and a subset of primary senescent cells is characterised by Notch

Since the secondary senescence clusters were only partially characterised by a paracrine senescence signature, we explored consistent differences between the secondary senescence and the primary OIS clusters. We first assessed the most differentially expressed genes and detected fibrillar collagens (Collagen 1A1, 3A1 and 5A2, Fig.2a). Downregulation of fibrillar collagens is consistently observed in senescence (*30*), but they failed to downregulate in the secondary senescence enriched cluster (Table 1, Fig.2a). A similar failure to downregulate collagens was recently reported in a more specialized primary senescence phenotype, induced by ectopic, temporal activation of Notch (*30*). The same report suggested that the secretome in RasV12−induced senescence was regulated by CCAAT/enhancer-binding protein beta (CEBPB), with Notch- induced senescence relying on transforming growth factor beta (TGFB) signalling (Fig.2b)(*30*). Several lines of evidence identify a NIS signature in the secondary senescence cluster. Firstly, IPA pathway analysis identifies TGFB1 as exclusively activated in the secondary senescence clusters compared to growing or the primary OIS clusters (Fig.2c). In contrast, RELA and IL1B pathways, regulators of the CEBPB transcriptome, were differentially activated in the primary OIS cluster (Fig.2c). Consistent with our RasV12 annotation, HRAS was exclusively activated in the primary OIS clusters (Fig.2c, Supplementary Fig.2a). Secondly, we profiled candidate genes involved in Notch signalling and TGFB activation. When plotting TGFB induced transcript 1, (TGFB1I1) with Notch-target connective tissue growth factor (CTGF) and CEBPB, we identified a significant (p<0.05) upregulation of CTGF and TGFB1I1 genes in the secondary senescence cluster with a simultaneous downregulation of CEBPB, significant on the protein, but not mRNA level (Fig.2d, e, p=0.016), resembling the TGFB/CEBPB bias in NIS. This bias was confirmed by qPCR on bulk sorted cells (Fig.2d, TGFB1 p=0.02, TGFBI p=0.05).

**Fig. 2.**
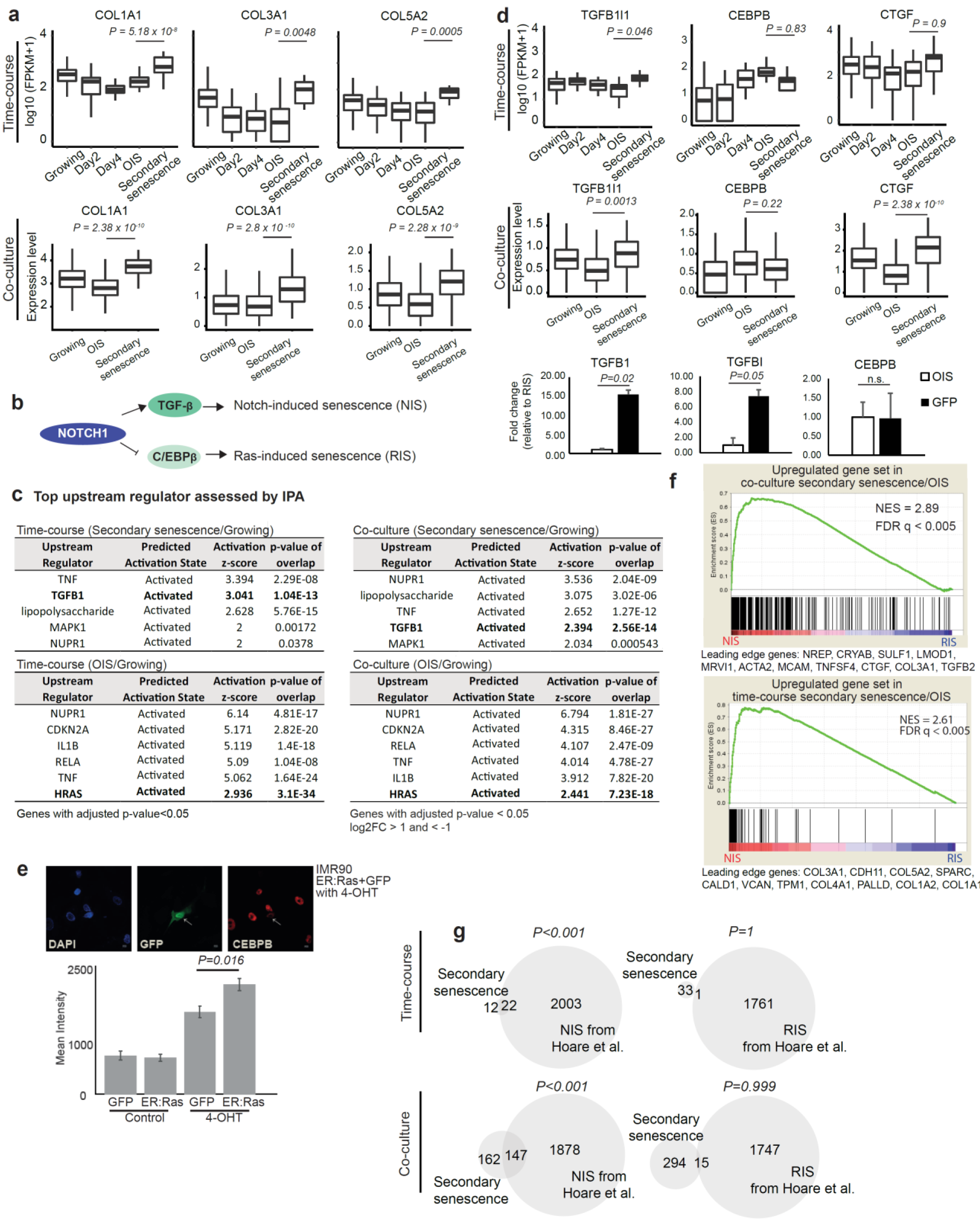
Secondary senescence comprises NIS signature in majority of cells. **a**. Box plots showing the expression of extracellular matrix organization genes *COL1A1*, *COL3A1* and *COL5A2* in the time-course and co-culture experiments (p-values < 0.05 as assessed by SCDE). The top and bottom bounds of the boxplots correspond to the 75 and 25th percentile, respectively. **b.** Model suggesting NIS and RIS are regulated by Notch1 through TGF-β and C/EBPβ respectively. **c.** IPA analysis of the two senescence clusters from the time-course and co-culture experiments relative to growing. **d.** Box plots for the expression of TGFB1I1, CTGF and CEBPB genes in the time-course (top) and co-culture experiments (middle). The top and bottom bounds of the boxplot correspond to the 75 and 25th percentile, respectively. Bar graphs denoting expression of *Tgfb1* (n=6), *TgfbI* (n=6), and *Cebpb* (n=3) mRNA as measured by qPCR in OIS and GFP cells (bottom) (TGFB1: t = −3.2317, df = 5.5117, p= 0.02; TGFBI: t = −2.2567, df = 9.8141, p=0.05; CEBPB: t = 0.068192, df = 3.2294, p=0.95 using unpaired Student’s t-test. Error bars represent SEM). **e.** A representative image of GFP (secondary senescence) and CEBPB (red) immunofluorescence in the co-culture experiment. Mean intensity for primary (ER:Ras) and secondary senescent cells (GFP) was measured (p=0.016 using unpaired Student’s t-test). Error bars are displayed as SEM. **f.** GSEA plots to assess the enrichment of secondary and primary OIS DE genes (from the time-course and co-culture experiments) in Hoare et al.’s NIS and RIS log2FC preranked genes. Normalised enrichment score (NES) and false discovery rate (FDR) are shown. **g.** Venn diagrams overlapping expression signatures from time-course (top panel) and co-culture (bottom panel) with NIS signature genes. (Secondary senescence: Secondary senescence/OIS upregulated genes; NIS: Hoare et al.’s NIS/RIS upregulated genes; RIS: Hoare et al.’s RIS/NIS upregulated genes).

Thirdly, we moved from a candidate gene approach to unbiased genome wide analysis. We calculated the enrichment of NIS and Ras induced senescence (RIS) signatures in the primary OIS and secondary senescence transcriptomes using gene set enrichment analysis (*31*) (GSEA) on ranked transcriptome differences between NIS and RIS (Fig.2f). Secondary senescence signatures from the time course and co-culture experiments were highly enriched in NIS (NES=2.61, FDR<0.005 for time course, NES=2.89, FDR<0.005 Fig. 2f). Primary OIS transcriptomes showed an enrichment for RIS (Supplementary Fig.2b). Finally, we interrogated the extent of NIS in the secondary senescence clusters by comparing the most differentially regulated genes (adjusted p<0.05) between RIS and NIS. We found a significant enrichment of NIS genes in our secondary senescence transcriptome in the time course and co-culture experiments with primary OIS signature being enriched for RIS genes (Fig.2f,g and Supp Fig.2b and 2c). In summary, our data identify a pronounced NIS transcriptional signature in secondary senescence and in a subset of primary senescent cells as an alternative endpoint to OIS.

### NIS is a secondary senescence effector mechanism during OIS

We next established Notch signalling as an effector mechanism in secondary senescence. We generated IMR90 fibroblasts with compromised Notch signalling by introducing a dominant negative form of mastermind like protein 1 fused to m-Venus (mVenus:dnMAML1) or empty vector (mVenus:EV) control and co-cultured with ER:Ras IMR90 cells (Fig.3a). At day 7 co-culture, mVenus:dnMAML1 compared to mVenus:EV cells exhibited lower expression of extracellular matrix gene *COL3A1* (p=0.02) and Notch target *CTGF* (p=0.056, Supplementary Fig.3a) as measured by qPCR, confirming impaired Notch signalling. Several lines of evidence show causal involvement of Notch signalling in secondary senescence. Firstly, we scored mVenus (YFP) signal between Notch perturbed (mVenus:dnMAML1) and mVenus:EV cells at day 0 (Growing) and day 7 tamoxifen co-culture with ER:Ras. At day 7, we observed significantly more mVenus:dnMAML1 compared to mVenus:EV cells (p=0.01), suggesting that primary OIS cells have less secondary senescence effect on neighbouring cells when harbouring perturbed Notch signalling (Supplementary Fig.3b). No significant difference in mVenus positive cells was observed in growing mVenus:EV compared to mVenus:dnMAML1 cells (p=0.38), showing that the dnMAML1 itself does not have an effect on cell number (Supplementary Fig.3b). Secondly, we scored EdU incorporation between mVenus:dnMAML1 and mVenus:EV cells at days 0 and 7 tamoxifen (Fig.3b). At day 7, we observed significantly more EdU incorporation in mVenus:dnMAML1 compared to mVenus:EV cells (p=0.01), with day7 mVenus:dnMAML1 cells showing similarly high levels of EdU incorporation as in growing mVenus:dnMAML1 and growing mVenus negative ER:Ras conditions (p=0.997 and p=0.08), suggesting that the induction of secondary senescence was abolished due to Notch perturbation (Fig.3b). As expected, ER:Ras cells showed low levels of EdU incorporation at day 7 tamoxifen (p=0.01 for ER:Ras with mVenus:dnMAML1 co-culture and p=0.0005 for ER:Ras with mVenus:EV co-culture, Fig.3b). Thirdly, we investigated SAHF in primary OIS and secondary senescence and showed that primary OIS cells displayed SAHF as expected (p=4.437×10^−6^, Supplementary Fig.3c). Secondary senescent cells (mVenus:EV) did not show significant SAHF formation when compared to OIS (p=0.32, Supplementary Fig.3c). This is consistent with previously published data in which impaired Notch signalling partially suppresses SAHF formation in primary senescence context (*32*). In summary, we show that Notch signalling mediates secondary senescence *in vitro*.

**Fig. 3.**
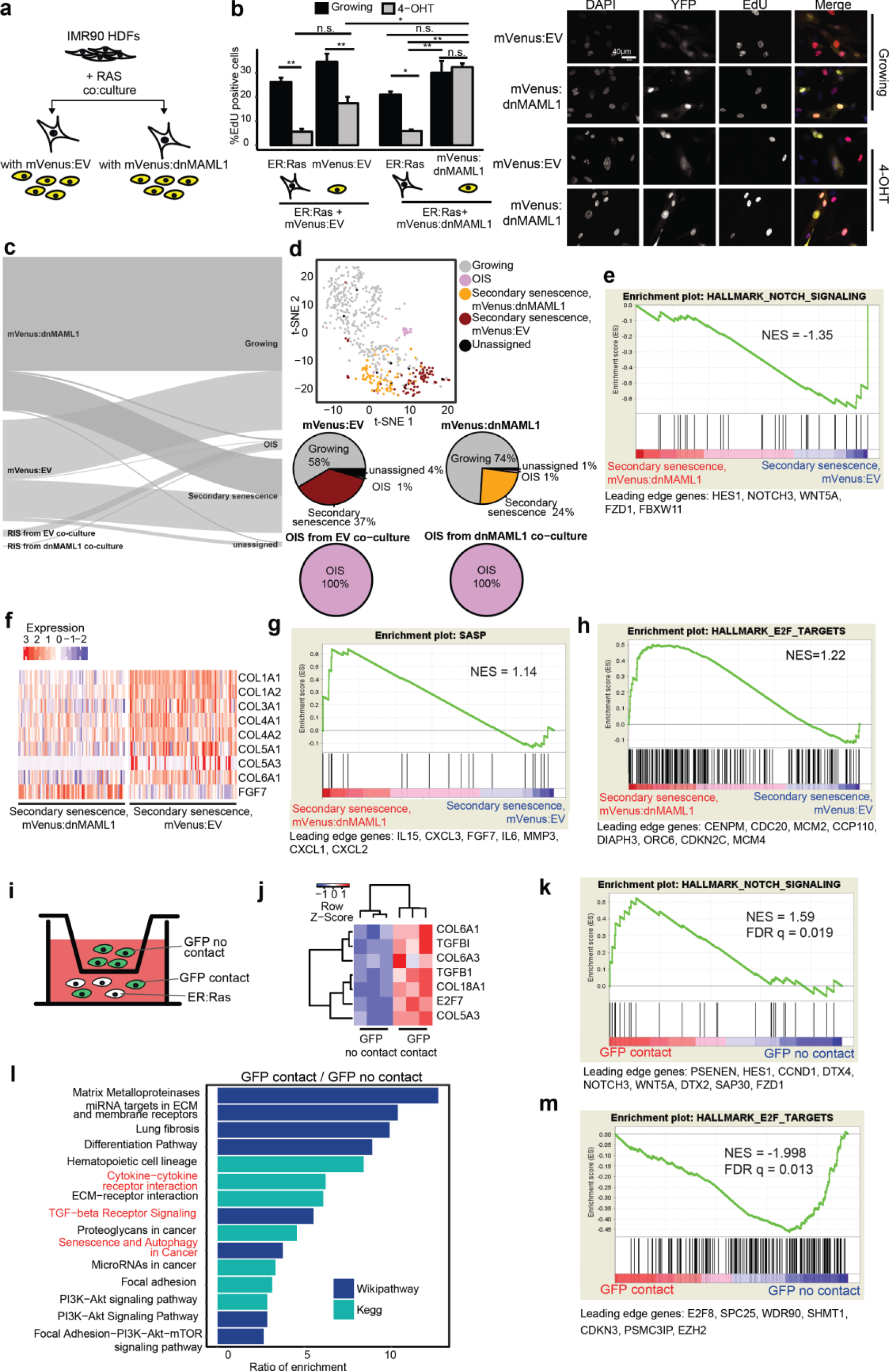
NIS mediates secondary senescence in vitro. **a.** Schematic representation of ER:Ras cells and mVenus:dnMAML1 or mVenus:empty vector (EV) cells co-culture experimental design. **b.** Bar plot showing EdU incorporation in growing (black) or senescent (grey) EV or dnMAML1 cells co-cultured with ER:Ras as proportion of all cells scored. Error bars are displayed as SEM; F[7,16] = 20.63, p<0.001 using one-way ANOVA with Tukey’s test. (n=3 for each condition). Representative images are shown on the right. **c.** Scmap-cluster projection of the dnMAML1 and EV 10x scRNA-seq dataset to the GFP co-culture 10x dataset (see Fig.1h). **d.** t-SNE plot of the single cells colored by the projection towards the GFP co-culture 10x dataset (see Fig.1h). The percentage of cells in each assignment was shown in pie charts. **e.** GSEA pre-ranked test showed the enrichment of Notch signaling in mVenus:EV identified as secondary senescence using scmap. **f.** Heatmap of single cell data comparing mVenus:EV and mVenus:dnMaml1 for collagens and SASP genes. Red despicts upregulated and blue downregulated. **g.** GSEA pre-ranked test showed the enrichment of SASP genes in mVenus:dnMAML1 identified as secondary senescence using scmap. **h.** GSEA pre-ranked test showed the enrichment of E2F targets in mVenus:dnMAML1 identified as secondary senescence using scmap. **i.** Schematic representation of the transwell co-culture assay of OIS cells and GFP cells. **j.** Heatmap of significantly differentially expressed genes (p<0.05) between GFP contact and GFP no contact cells obtained from RNA-seq. **k.** GSEA pre-ranked analysis showed the significant enrichment of Notch signaling in GFP contact cells in comparison to GFP no contact cells. **l.** Ratio of enrichment of over-representation pathway analysis for GFP contact/GFP no contact differentially expressed genes (p<0.05). **m.** GSEA pre-ranked analysis showed the significant enrichment of E2F targets in GFP no contact cells in comparison to GFP contact cells. Leading edge genes are indicated below the plot.

To establish transcriptional differences between secondary senescence with and without Notch signalling, we generated single cell transcriptomes from IMR90 mVenus:EV and mVenus:dnMAML1 co-cultures with ER:Ras IMR90 at day 7. To integrate the new data set with our previous secondary senescence transcriptomes (Fig.1h growing, OIS and secondary senescence), we projected the mVenus:EV and mVenus:dnMAML1 using sc-map (*33*).

Sc-map clearly matches all primary senescent cells containing RasV12 to the OIS population (Fig.3c). Sc-map identifies significantly more secondary senescence cells in mVenus:EV compared to mVenus:dnMaml1 (Fig.3c, 37% vs 24%, Chi-squared test, p=0.00062), confirming a role of Notch in secondary senescence mediation. To explore transcriptomic differences between secondary senescence, we plotted all cells using Seurat, which separated mVenus:EV and mVenus:dnMAML1 into distinct secondary senescence clusters (Fig.3d). We confirmed differences in the activation of Notch pathway between mVenus:EV and mVenus:dnMaml1 by GSEA analysis (Fig.3e, NES=-1.35) and on the gene level for fibrillar collagens (Fig.3f, p<0.05). Interestingly, Notch signalling seems to blunt the cytokine response in senescence as SASP factors (Fig.3g, NES=1.1) and the interferon gamma response (Supp.Fig.3d, NES=1.48) are differentially regulated between mVenus:EV and mVenus:dnMaml1 as judged by GSEA. Importantly, E2F targets, whose downregulation is one of the hallmarks of senescence, are upregulated in mVenus:dnMaml1 cells compared to mVenus:EV (Fig. 3h, p=not significant), which offers an explanation for the strong phenotype differences we observed between the two conditions (see Fig.3b).

Notch induces senescence in a juxtacrine manner through cell-to-cell contact. Therefore, we performed transwell experiments to verify the effect of cell-to-cell contact on the secondary senescence transcriptome. We co-cultured ER:Ras cells with GFP cells in a twelve well plate (GFP contact, Fig. 3i) and GFP cells on their own in the transwell of the same well (GFP no contact). In this setting, GFP no contact cells shared media with ER:Ras fibroblasts, where cytokines can be transferred, but no cell-to-cell contact is possible. We performed bulk RNA-sequencing of GFP contact and no contact fibroblasts 7 days after tamoxifen induction and confirmed enhanced expression of previously observed marker genes for NIS secondary senescence in GFP contact cells (Fig.3j). In addition, GSEA confirmed enrichment of Notch (NES=1.59, FDR q=0.019) and TGF beta (NES=1.87, FDR q=0.0016) signalling (Fig.3k and Supplementary Fig.3e) in GFP contact cells. Pathway analysis confirmed significant upregulation of previously described senescence pathways such as “Senescence and Autophagy in Cancer”, “Matrix Metalloproteases” and “PI3K-AKT-mTOR” in GFP contact compared to GFP no contact cells (Fig.3l). Equally, GSEA showed repression of E2F target genes in GFP contact compared to GFP no contact fibroblasts (Fig.3m) except for E2F7, which is known to be upregulated in senescence (Fig.3j). GSEA analysis suggests that the global differences between GFP contact and no contact cells resemble the differences between mVenus:EV and mVenus:dnMaml1 secondary senescence (Supplementary Fig.3f).

OIS induction is a multi-step process with an early proliferative phase at days 1-3, followed by a phenotype transition phase at days 3-5, leading to established senescence from day 7 after RasV12 expression (*24*). To compare the impact of the different phases of primary OIS induction onto secondary senescence, we co-cultured mVenus:EV or mVenus:dnMAML1 cells repeatedly with ER:Ras cells at days 3-6 or at days 7-10 after RasV12 induction for a total of 4 cycles (Suppl.Fig.3g). As expected, ER:Ras cells showed low levels of EdU incorporation in mVenus:dnMAML1 (Day3 and Day7 p<0.001) co-culture or mVenus:EV co-culture (Day3 and Day7 p<0.001) (Supp. Fig.3h,i) as a result of primary OIS. Co-culturing mVenus:EV cells with ER:Ras cells in transition phase (days 3-5 after RasV12 induction), lead to a significant reduction in EdU when compared to uninduced co-cultures (p<0.001, Supp. Fig.3h), suggesting that secondary senescence was induced by transition phase primary OIS cells. The transition phase effect is Notch dependent since it cannot be induced in mVenus:dnMAML1 cells (p=0.12, Supp. Fig.3i). In contrast, by co-culturing mVenus:EV cells with primary OIS cells in established senescence phase (7-10 after RasV12 induction), we were unable to detect a reduction in EdU incorporation in mVenus:EV cells compared to uninduced co-cultures (p=0.59, Supp. Fig.3h), mirroring results obtained in mVenus:dnMAML1 co-cultures (p=0.99, Supp. Fig.3i). Moreover, from day 4 co-culture, we detected a significant upregulation of Notch1 on mVenus:EV (p= 0.041 day4, p=0.038 day) and mVenus:dnMaml1 (p=0.023 day, p=0.046 day7) cells compared to growing, providing a pathway to NIS induction (Supp. Fig.3j). These results highlight a need for ER:Ras fibroblasts to be in phenotype transition phase to mediate secondary senescence via Notch1. Overall, our data identify Notch as a key mediator of secondary senescence.

## Secondary senescent hepatocytes are characterised by NIS signature

To test the involvement of NIS *in vivo*, we utilised a model where primary senescence is induced in a subpopulation of hepatocytes following *Mdm2* deletion (*34*). This model is activated by hepatocyte-targeted recombination of *Mdm2* (βNF induction AhCre, Mdm2−), resulting in primary senescence in Mdm2- cells. Mdm2- hepatocytes induce secondary senescence in local hepatocytes^33^ (Fig.4a). In this model, the presence of p53 induction through Mdm2 deletion with medium levels of CDKN1A (non senescence/primary p<0.001) marks primary senescence induction (*34*) (Fig.4b, Supplementary Fig.4a,b). Physiological levels of p53 and high levels of CDKN1A (CDKN1A expression secondary/primary p<0.0001) marks secondary senescence in Mdm2 normal (Mdm2+) hepatocytes as described (*34*) (Fig. 4b). Based on these characteristics, cells can be readily distinguished by immunohistochemistry with 23% of primary and 10% of secondary senescence hepatocytes detected (Supplementary Fig.4a). We have previously shown that both subpopulations of hepatocytes upregulate senescence markers (gH2AX, IL1A, SA-beta Galactosidase) and reduce BrdU incorporation (*34*).

**Fig. 4.**
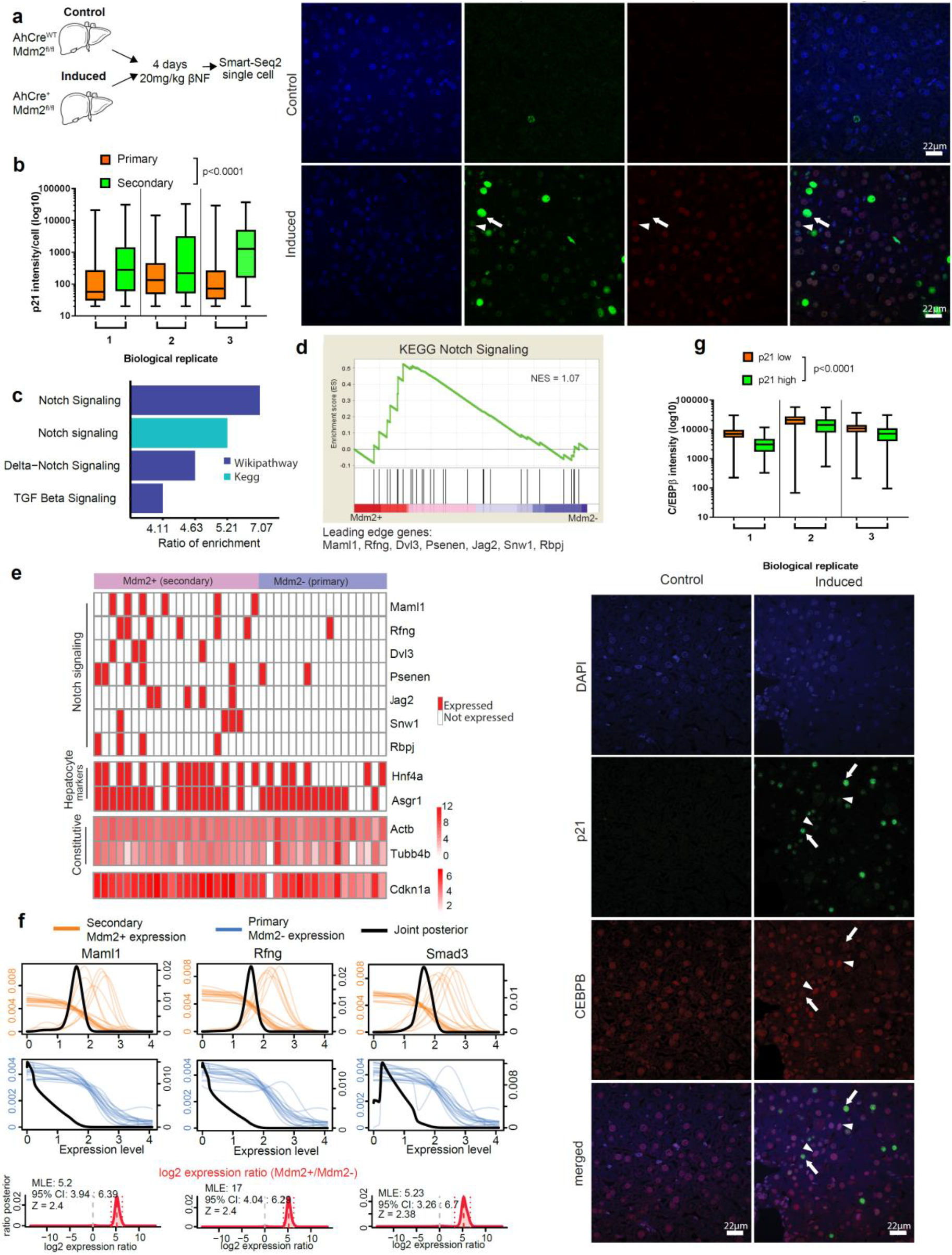
Notch signaling mediates secondary senescence *in vivo*. **a**. Schematic representation of *in vivo* single-cell experiment. **b.** Representative immunofluorescence images of liver section from induced AhCre+Mdm2fl/fl and control AhCre^WT^Mdm2^fl/fl^ mice. The accumulation of p53 through Mdm2 deletion with low levels of p21 marks senescent cells that are induced intrinsically (arrowhead), whereas physiological levels of p53 and high levels of p21 marks secondary senescent cells in Mdm2 normal hepatocytes (arrow). The boxplot showed p21 intensity in primary versus secondary senescent cells. (senescence: F[1,50291] = 2766, p<0.0001; biological replicates: F[2,50291]=283.2, p<0.0001; senescence x biological replicates: F[2,50291]=280.5, p<0.0001 using two-way ANOVA). Scale bar 22μm. **c.** Ratio of enrichment for over-representation pathway analysis for Mdm2+ (secondary) genes. **d.** GSEA revealed Notch signaling as one of the top enriched pathways in Mdm2+/Mdm2- cells (Normalised enrichment score, NES=1.07). Leading edge genes are indicated. **e.** Gene expressions for Notch signaling pathway, hepatocyte markers and *Cdkn1a* gene are plotted in Mdm2+ and Mdm2- cells. Constitutive genes and *Cdkn1a* were colored by their expression relative levels (binary: red expressed, white not expressed). **f.** SCDE expression for Maml1, Rfng and Smad3 in Mdm2+ cells (orange lines) and Mdm2- cells (blue lines). Joint posterior is marked by black line. Fold change of the genes in Mdm2+/Mdm2- is indicated in red and dotted lines mark the 95% confidence interval. MLE: Maximum likelihood estimation; CI: Confidence interval; Z: z-score. **g.** Representative immunofluorescence images of liver section from induced and control mice. Primary senescent cells showed low p21 level and high CEBPB level (arrowheads), whereas secondary senescent cells showed high p21 and low CEBPB levels (arrows) (p21: F[1,60145] =353.3, p<0.0001; biological replicates: F[2,60145]=1044, p<0.0001; p21 x biological replicates: F[2,60145]=8.96, p<0.0001 using two-way ANOVA). Scale bar 22μm.

To establish if primary and secondary senescence can be distinguished based on the transcriptome *in vivo*, we performed single-cell RNA-seq on hepatocytes using Smart-seq2 (Fig.4a). After filtering (Supplementary Fig.S4b,c and Supplementary Table 1), we retained 39 single cells from induced Mdm2 deleted mouse liver for downstream analysis. We distinguished Mdm2- cells from Mdm2+ hepatocytes by the absence of mapping reads over exon 5 and 6 of the Mdm2 gene (Supplementary Fig.4d). We detected expression of CDKN1A in both senescent populations consistently with the differences in CDKN1A protein levels detected by immunohistochemistry (Fig.4b), with lower (but not significant) CDKN1A expression in Mdm2- compared to Mdm2+ hepatocytes (Supplementary Fig.4e), enabling us to distinguish primary and secondary senescence. To verify a senescence phenotype in both Mdm2- and Mdm2+ hepatocyte populations, we conducted pathway analysis with upregulated pathways being enriched in p53 signalling pathways, including CDKN1A, DNA damage response and cytokine signalling (Supplementary Fig.4f). We next asked if NIS plays a role in secondary senescence *in vivo* analysing our single cell data using three independent methods. Differential expressed genes between Mdm2+ and Mdm2- cells were identified using SCDE (Table 1) and genes were ranked between Mdm2+/Mdm2- cells for downstream analysis. Firstly, pathway analysis revealed an enrichment in Notch signalling (Ratio of enrichment (RE) 7.07), Delta-Notch signalling (RE 4.63) and TGFB (RE 4.11) signalling pathways (Fig.4c). Secondly, GSEA revealed Notch signalling pathway (NES=1.07) as one of the top 20 Kegg pathways enriched in Mdm2+/Mdm2- (Fig.4d) with leading edge genes MAML1 and JAG2 detectable mainly in the Mdm2+ cells (Fisher’s exact test=6.93×10^−7^, Fig. 4e). Housekeeping and hepatocyte specific genes were expressed to the same level in the majority of cells regardless of Mdm2 status (Fig.4e). Thirdly, SCDE analysis confirmed the specific upregulation of Notch and TGFB targets MAML1 (aZ=0.4) and RFNG (aZ=0.39) with effector protein SMAD3 (aZ=0.26) in Mdm2+ compared to Mdm2- hepatocytes (Fig.4f). To assess the proposed TGFB/CEBPB bias between primary and secondary senescence *in vivo*, we stained livers from uninduced and induced mice for CDKN1A and CEBPB by immunohistochemistry. Consistent with our *in vitro* data, we observed significantly higher CEBPB protein in primary (p<0.0001, Fig. 4g) compared to secondary senescent hepatocytes. These lines of evidence show that secondary senescent hepatocytes are characterised by a NIS signature *in vivo.*

## Discussion

Cancer heterogeneity is an expanding field of research. Much less is known about cellular heterogeneity in a pre-cancerous state. Are all cells reacting similarly to oncogene activation or does an oncogenic insult result in a heterogeneous population? Understanding heterogeneity in a pre-cancerous state will inform distinct propensities for transformation in subpopulations. Our study uncovers heterogeneity in primary OIS and secondary senescence transcriptomes following an oncogenic insult using single cell approaches.

Paracrine induction of senescence is thought to be the main mediator of secondary senescence in OIS (*16,22*). Our results challenge this canonical view implicating NIS as synergistic driver of secondary senescence *in vitro,* in the most studied OIS background (RasV12) and in the liver *in vivo*.

Currently, primary and secondary senescent cells are not thought of as functionally distinct endpoints. However, we provide strong evidence for differences between primary OIS and Notch-mediated juxtacrine secondary senescence as they display distinct gene expression profiles and potentially different transformation potential (*22,30*). Some of our findings point to a functional diversification, for example the blunted SASP response and the induction of fibrillar collagens in secondary senescence compared to OIS. It will be important to understand the role of secondary NIS mediated senescence in cancer etiology: Why do these neighbouring cells have to senesce? How are they contributing to the micro-environment and malignancy? In one scenario, the secondary senescence response is a failsafe mechanism to prevent cells potentially carrying a genetic lesion from avoiding senescence. In the other scenario, the mechanism of secondary senescence induction might affect the microenvironment and malignancy differently. Therefore, heterogeneity in senescence induction has implications therapeutically, especially in the application of senolytic drugs.

We identified two transcriptional end-points for primary OIS, namely a Ras driven and a NIS programme. Notch signalling is mediated through cell-to-cell contact(juxtacrine), and Hoare and colleagues (2016) have shown that it can be a transient state towards primary senescence induction (*30*). However, our data indicate cells carrying a composite transcriptional signature of paracrine and juxtacrine events as a facultative end-point for cells with detectable Ras activation (primary). Within the Ras activating cells, the minority progresses to the NIS resembling state. This transcriptional heterogeneity within pre-cancerous populations is an important concept, and the transformation potential of these heterogeneous populations will need to be addressed in the future.

## Supporting information

compiled supplement

## Acknowledgments

**Funding:** K.K. was supported by the Wellcome Trust (grant number 105641/Z/14/Z). T.C. is supported by a Chancellor’s Fellowship held at the University of Edinburgh. TGB is funded by the Wellcome Trust (WT107492Z). Research in the N.N. lab was partially supported by IDeA grant P20GM109035 (Center for Computational Biology of Human Disease) from NIH NIGMS and grant 1R01AG050582-01A1 from NIH NIA to NN. N.P was supported by a Ph.D studentship funded by the Wellcome Trust Sanger Institute and the Royal Thai Government. P.D.W. was funded by P01 AG031862. Work in the Green laboratory was supported by Bloodwise, Cancer Research UK, Wellcome Trust, the National Institute for Health Research Cambridge Biomedical Research Centre, the Cambridge Experimental Cancer Medicine Centre, and the Leukemia & Lymphoma Society of America. J.-C.A. was supported by a CRUK-Career Development Fellowship C47559/A16243. We would like to thank the CRUK Beatson Histology core facility for help with immunohistochemistry. We would like to thank Prof. Chris Ponting and Dr. Andy Fynch for critical reading of the manuscript.

## Author contributions

T.C., K.K. and N.N. conceived the study. Y.V.T. performed all bioinformatics analysis. N.P. performed the majority of the experiments and data analysis with help from A.S., A.Q., N.T. and K.K. In vivo work and data analysis was carried out by C.K. M.M. and T.G.B. P.D.A., J.-C.A. and A.R.G. gave feedback on the study and commented on the manuscript. K.K., T.C. and Y.V.T. wrote the manuscript.

## Data Availability

The sequencing data of Smart-seq2 time-course single-cell RNA-seq experiment, 10x co-culture single-cell 3’ expression experiment, and Smart-seq2 mouse *in vivo* liver hepatocyte single-cell RNA-seq are accessible through GEO accession number GSE115301. All imaging data are available as Mendeley data set under doi:10.17632/y76pb7s8h3.1.

## Supplementary Materials List

Materials and Methods

Figures S1-S4

Tables S1-S4

**Table 1. Differential expression analysis of all transcriptome data sets. Comparisons are indicated in the header of each subtable.**

