## Supplementary material for "Notch signalling mediates secondary senescence": compiled supplement

**This PDF file includes:**

Materials and Methods
Figs. S1 to S4
Tables S1 to S4 submitted as separate files

### Materials and Methods

#### Tissue culture

ER:Ras$^{G12V}$ -expressing cells were maintained and senescence induced as described *(24)*. ER:IMR90 cells were co-cultured IMR90:GFP, empty vector fused with mVenus (EV:mVenus) or with a dominant negative form of MAML1 fused with mVenus (dnMAML1:mVenus) cells at 1:10 ratio.

#### Transwell assay

ER:Ras$^{G12V}$-expressing cells were co-cultured with IMR90:GFP. The co-cultured cells were placed into the lower chamber of a transwell system (density $5x10^3$ cells/well). Another pure population of IMR90:GFP cells were cultured in the upper chamber of the transwell system. All cells were maintained in 4-hydroxytamoxifen for 7 days.

#### Single cell data generation

Smart-Seq2 was performed on sorted ER:IMR90 cells or hepatocytes as previously described *(37)*. Single cell data for all co-culture experiments were generated using the 10X Chromium platform, following the manufacturer's instructions.

#### Flow cytometry

Flow cytometry was performed as previously described (*37)* using 7-AAD, DAPI, anti-Notch1-PE (R&D systems, FAB5317P, 1:20), anti-JAGGED1-APC (R&D systems, FAB1726A, 1:20) where indicated using BD FACSAria II (BD Biosciences, San Jose, CA) and the BD CellQuest PRO software (BD Biosciences, San Jose, CA). Flow data were analysed with FlowJo v10 (Tree Star, Ashland, OR).

#### RNA extraction

RNA was extracted using the Qiagen Mini Kit. All RNA passed with a RIN of 9 or above as determined by Bioanalyser profiling. Ribosome depletion was performed prior to bulk RNA sequencing.

**qPCR**

qPCR was performed on a LightCycler 480 (Roche) using Sybr Green method as previously described[9]. Delta delta Ct method was used for quantification with error bars resulting from the delta Ct expression of replicates. A two-sided t-test was used to calculate p-values.

**EdU incorporation and SA-beta Gal staining**

EdU incorporation and SA-beta gal staining was performed as previously described (*9)*. EdU incorporation was detected using the Click-iT™ EdU Alexa Fluor 555 imaging kit (Invitrogen/Molecular Probes, Eugene, OR). For stable cell cycle arrest, cells were co-cultured for two weeks, separated by FACS and cultured as OIS and GFP cells for another week before pulsing them with EdU for 24 hours.

**Confocal microscopy and Image analysis**

BriteMac confocal microscope was used to visualize cells at 40x. Images were analysed using ImageJ. Percentages of SAHF, YFP/GFP and EdU-positive cells were calculated by assessing 1600-2000 cells (triplicate counts) per experiment.

**Animal models**

Animal welfare conditions have been previously described (*38*). All animal experiments were carried out under procedural guidelines, severity protocols and within the UK with ethical permission from the Animal Welfare and Ethical Review Body (AWERB) and the Home Office (UK). AhCre+/WT *Mdm2*fl/fl and AhCreWT/WT *Mdm2*fl/fl mice (colony N4 C57/Bl6J background) were crossed and male mice of 10-16 weeks used in experiments. Genotyping and i.p. injection of β-Napthoflavone (βNF, Sigma UK) at 20mg/kg were performed as previously described (*34*). Livers were harvested and partially stored in paraffin blocks following fixation in 10% formalin (in PBS) for 18 hours prior to embedding.

**Hepatocyte isolation**

*Ex vivo* primary hepatocytes were isolated using a modified retrograde perfusion technique as previously described (*34*). Hepatocytes were purified by pelleting through a 40% (v:v) percoll gradient prior to FACS sorting.

**Immunohistochemistry**

Immunohistochemistry was performed as described[33]. Three µm thick paraffin sections were double stained for p53/CDKN1A and CDKN1A/CEBPb using the CDKN1A clone HUGO291H (a gift from Serrano lab, CNIO in Madrid), C/EBPb clone 1H7 (Abcam) and NCL-L-p53-CM5p (Leica Biosystems). Detection was performed with TSA-Cy3 (Perkin Elmer, NEL744B001KT, 1:50) and TSA-FITC (Perkin Elmer, NEL741B001KT, 1:50). Images were captured on a Zeiss 710 Upright Confocal Z6008 microscope. Stained slides were scanned using the Opera Phoenix High Content screening system (Perkin Elmer) scanner and analysed using the Columbus software.

**Bioinformatics analysis**

**Sequencing reads processing, alignment and quantification of time-course experiment**

Smart-Seq2 generated paired-end reads were quality trimmed using Trim galore (http://www.bioinformatics.babraham.ac.uk/projects/trim_galore/) and aligned to the human reference genome, hg19, neomycin sequence from pLNCX2-ER-ras_neo, ERCC spike-in sequences and RasV12 using HISAT v2.0.1beta (*39*). Cells with less than 200,000 hg19 aligned reads, and a ratio of ERCC RNA spike-in control aligned reads to total aligned reads greater than 0.5 were omitted. hg19 aligned reads were randomly downsampled to 200,000 reads. Genes were quantified using HTSeq-0.6.1(*40*). Cells with more than 80,000 total gene counts and at least 500 genes with at least one count were used for downstream analysis. 224 IMR90 cells (100 Growing cells, 41 Day 2 cells, 42 Day 4 cells and 41 senescent cells) passed this second filtering step and used for downstream analyses.

**Sequencing reads processing, alignment, quantification and analysis of 10x Chromium RNA-seq data**

Cell Ranger 2.0.1 (10x Genomics) was used to align the GFP and ER:Ras$^{G12V}$ co-culture 10x Chromium RNA-seq reads to hg19, TurboGFP, puromycin sequence from pGIPZ and neomycin sequence from pLNCX2-ER-ras_neo, and to generate gene-cell matrices. The growing and senescence dataset were aggregated using "cellranger aggr". The data were subsequently processed using Seurat 2.3.0 with cells with less than 15% mitochondrial reads and at least 2500 number of genes being retained. Seurat 2.3.0 with the default parameters (unless otherwise stated) was used to generate the t-SNE plots (resolution:0.4; dimensions used: 1:15) and three clusters were identified using sparcl 1.0.3 (https://cran.r-project.org/web/packages/sparcl/index.html). SCDE v1.99.1 was used to identify differentially expressed genes between OIS cluster and secondary senescent cluster. The DE genes (p-values < 0.05) (Table 1) were used as the defined gene sets for GSEA Preranked analysis of NIS and RIS log2FC ranked genes. GFP+ cells were identified as cells with > 0.3 normalized expression of GFP or puromycin and Ras+ cells were identified as cells with non-zero expression of neomycin or one or more reads supporting the G>T mutation at Chr11:534288 as identified by FreeBayes v0.9.20-8-gfef284a(*41*). Integration analysis between Smart-seq2 time-point data and 10x data were performed using the canonical correlation analysis in Seurat 2.3.0, in which the union of the top 50 highest dispersion genes and the first two dimensions were used.

Cell Ranger 2.0.1 (10x Genomics) was used to align the 10x Chromium RNA-seq reads from mVenus:dnMAML1 or EV:dnMAML1 co-cultured with ER:Ras$^{G12V}$ cells to hg19, mVenus sequence, puromycin sequence from pLPC-puro and neomycin sequence from pLNCX2-ER-ras_neo to generate gene-cell matrices. mVenus cells were identified as cells with more than zero normalized expression of mVenus or puromycin and Ras+ cells were identified as cells with non-zero expression of neomycin or one or more reads supporting the G>T mutation at Chr11:534288 as identified by FreeBayes v0.9.20-8-gfef284a *(41)*. The data were subsequently processed using Seurat 2.3.0 with cells with less than 10% mitochondrial reads and at least 2500 number of genes being retained. Seurat 2.3.0 with the default parameters (unless otherwise stated) was used to generate the t-SNE plots (resolution:0.6; dimensions used: 1:7). The cells were projected to the 10x Chromium GFP and ER:Ras$^{G12V}$ co-culture dataset using scmap-cluster v1.4.1.

**Sequencing reads processing, alignment and quantification of *in vivo* data**

Smart-Seq2 generated paired-end reads were quality trimmed using Trim galore (http://www.bioinformatics.babraham.ac.uk/projects/trim_galore/) and aligned to the mouse reference genome mm10 and ERCC spike-in sequences using HISAT v2.0.1beta (*39*). The mm10 aligned reads were randomly downsampled to 50,000 reads. Cells with less than 50,000 reads, less than 20,000 gene count, less than 500 genes with at least one read detected and with the log-transformed number of expressed genes and library size of 3 median absolute deviation below the median value were removed (*42*). 39 single cells from the induced hepatocytes and 19 cells from the uninduced hepatocytes passed these filters. 22 primary senescent cells were identified from the induced hepatocytes as cells with no reads mapping over exon 5 and 6 (chr10:117695953-117696049, chr10:117696381-117696439, chr10:117701565-117701614 and chr10:117702202-117702335) of Mdm2 gene before the downsampling, and the remaining 17 expression between induced and uninduced hepatocytes, and upregulated genes (adjusted cells were classified as secondary hepatocytes. Differential genes expression between Mdm2+ cells and Mdm2- cells was identified using SCDE v1.99 and log2FC ranked gene list from SCDE was used in GSEA pre-ranked analysis. Genes with more than zero log-transformed normalized count[43] were labeled red, and otherwise white in the binary heatmap. Pathway enrichment was identified using WebGestalt (*43*) with genes that have a z-score of greater than 2 in Mdm2+ cells /Mdm2- comparison.

**Differential gene expression analysis and temporal ordering of cells**

We used raw counts from HTSeq-Count as an input to single-cell differential expression (SCDE v1.99.1) (*26*) for differential gene expression analysis between growing and senescence. Cut-off for significantly differentially expressed (DE) was set at 0.05. The expression magnitude (fragments per million) was obtained from SCDE and converted to FPKM as an input for Monocle2 (*27*). Monocle2 was used to order the transitions of senescent cells of different time points at a pseudo-temporal resolution, and single-cell data were reduced to a 2-dimensional space by using the DDRTree algorithm implemented in Monocle2 (*26*). Specifically, DE genes between senescence and growing conditions that were identified in SCDE were used to define the trajectory. A consensus clustering approach, SC3, was also applied to the raw count of single cells and used to cluster senescent cells.

**Detection of Ras$^{V12}$ construct in Smart-seq2 dataset**

We counted reads with a G>T mutation at Chr11:534288 using samtools/1.2 mpileup and bcftools/1.2 (*44*). Cells with more than 1 read supporting over G>T mutation or at least 9 reads mapping to the neomycin sequence are considered as Ras$^{V12}$ positive cells.

**Paracrine-induced senescence and RIS microarray data analysis**

Log2 RMA signal intensity of RIS IMR90 cells and IMR90 co-cultured in transwells with RIS cells were obtained from GEO GSE41318. Differentially expressed genes were identified using LIMMA and an adjusted p-value of 0.05 was used as the cut-off for significant genes.

**Notch and Ras-induced senescence data and GSEA analysis**

We used NIS and RIS RNA-seq data with accession number GSE72404. Reads were aligned to as described above. Differential gene expression analysis between NIS and RIS was performed using DESeq2 (*46*). The log2 fold change for each gene was used to rank the list of genes in GSEA Preranked analysis (*31*). Differentially expressed (DE) genes between senescence top and bottom were identified using SCDE with a p-value cutoff of 0.05. The DE genes defined the gene set in GSEA Preranked analysis.

**Sequencing reads alignment and quantification of transwell bulk RNA-sequencing data**

Reads were aligned to the human reference genome hg19 using HISAT v2.0.1beta (*39*) and those that mapped to annotated genes were quantified using HTSeq-0.6. (*40*). Differential gene expression was determined using DESeq2 v1.22.1 (*46*). Over-representation analysis was performed using WebGestalt (*43*) and GSEA pre-ranked analysis was performed using the ranking of genes based on the log2FC between GFP contact and GFP no contact.

**Statistical analysis**

All t-tests and one-way ANOVA for the *in vitro* data were performed in R. TukeyHSD was used as the post hoc test for one-way ANOVA. For the *in vitro* data, each experiment and measurement were performed in triplicates unless otherwise specified in the figure legends. Barplots are represented as means with SEM. Statistical significance was set at $p < 0.05$. T-test for the *in vivo* data was performed in R and the two-way ANOVA followed by Tukey's test for the *in vivo* data was performed using GraphPad Prism.

### Supplementary Fig. 1

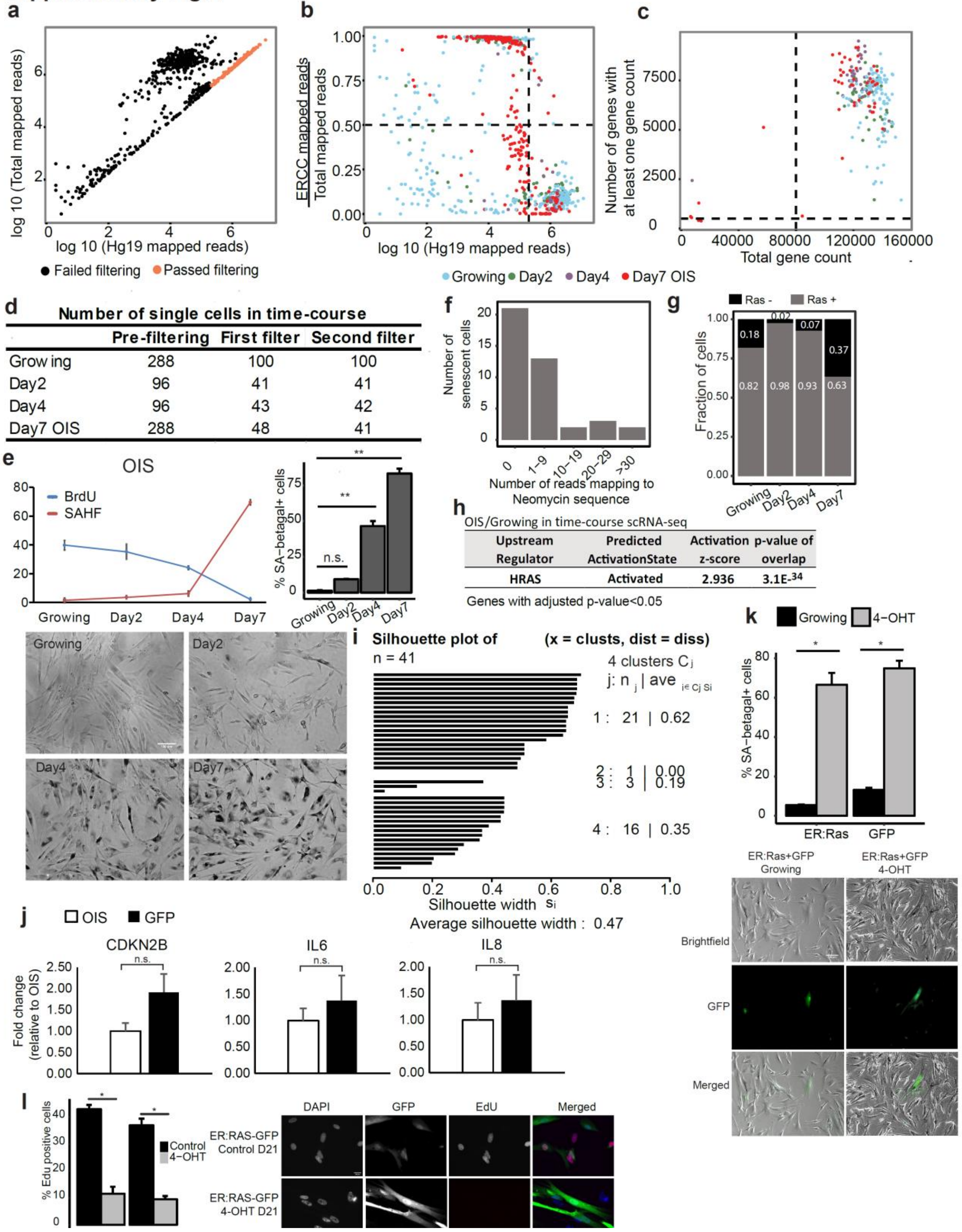


**d** Number of single cells in time-course

| | Pre-filtering | First filter | Second filter |
|---|---|---|---|
| Growing | 288 | 100 | 100 |
| Day2 | 96 | 41 | 41 |
| Day4 | 96 | 43 | 42 |
| Day7 OIS | 288 | 48 | 41 |

**h** OIS/Growing in time-course scRNA-seq

| Upstream Regulator | Predicted ActivationState | Activation z-score | p-value of overlap |
|---|---|---|---|
| HRAS | Activated | 2.936 | $3.1E^{-34}$ |

Genes with adjusted p-value<0.05

**Supplementary Fig. 1. Quality control and processing of single cells. a, b.** Filtering according to total mapped reads. Cells with less than 200,000 human aligned reads and with a ratio of ERCC RNA spike-in control aligned reads to total aligned reads that is greater than 0.5 were removed. The number of cells that passed this filtering step was shown in **d. c.** The second filtering step was performed to retain cells that have greater than 80,000 total gene counts and at least 500 genes with at least one count. Cell were then normalised by downsampling to 200,000 aligned reads for downstream analysis. **e.** BrdU, SAHF and SA-beta galactosidase counts in ER:Ras fibroblasts. Days indicated time of tamoxifen treatment. Error bars are SEM, n=3 for each time point. $F[3,8] = 234.8$, $p<0.001$; **$p < 0.001$ using one-way ANOVA with Tukey's test. Scale bar 100μm. **f.** Number of reads aligning to the neomycin sequence from the pLNCX2-ER-ras_neo construct insenescent single cells in the time-course experiment. **g.** Fraction of cells that are RasV12+ in eachcondition in the time-course experiment. **h.** IPA analysis of OIS/growing in time-course scRNA-seq. **i.** Silhouette plot to assess the quality of clustering. The average silhouette width was 0.47. **j.** Box plots for gene expression of *Cdkn2b* (n=3), *IL6* (n=3), and *IL8* (n=3) mRNA measured by qPCR in OIS and GFP cells. Unpaired Student's t-test showed no significant difference in senescent markers expression between OIS and GFP cells. Error bars represent SEM. **k.** SA-beta galactosidase counts in OIS and GFP cell s. (OIS t = 10.199, df = 2.0096, p= 0.009; GFP t = 15.239, df = 2.3673, p= 0.002 using unpaired Student's t-test). Scale bar 100μm. **l.** Bar plots showing EdU incorporation in GFP cells co-cultured with ER:Ras cells after 21 days as proportion of all cells scored. Error bars are displayed as SEM; **$p<0.001$, *$p<0.05$. Representative images are shown. Scale bar 20μm. (ER:Ras t = -9.899, df = 2.86 68, p= 0.0024; GFP t = 10.395, df = 3.3348, p= 0.0012 using unpaired Student's t-test) (n=3 for each).

### Supplementary Fig. 2

**a** **Top activated and shared upstream regulator assessed by IPA**

Time-course
(Secondary senescence / OIS)

| Upstream Regulator | Activation z-score | p-value of overlap |
|---|---|---|
| decitabine | 2.058 | 1.16E-13 |
| TGFB1 | 1.839 | 1.64E-12 |
| Brd4 | 0.788 | 2.55E-13 |
| TP53 | 0.653 | 1.44E-14 |
| ERBB2 | 0.611 | 2.01E-13 |
| forskolin | 0.488 | 4.7E-13 |
| D-glucose | 0.271 | 1.45E-12 |
| ERK | -0.193 | 1E-15 |
| KRAS | -0.805 | 1E-13 |
| EGF | -0.835 | 2.57E-16 |
| lipopolysaccharide | -0.97 | 1.01E-13 |
| HRAS | -3.116 | 1.09E-17 |

Co-culture
(Secondary senescence / OIS)

| Upstream Regulator | Activation z-score | p-value of overlap |
|---|---|---|
| PD98059 | 4.571 | 5.34E-29 |
| U0126 | 3.069 | 5.91E-29 |
| dexamethasone | 2.646 | 2.65E-33 |
| TGFB1 | 2.622 | 1.3E-48 |
| MYCN | -2.054 | 1.07E-22 |
| Cg | -2.253 | 2.03E-24 |
| EGF | -2.322 | 4.52E-22 |
| EGFR | -2.617 | 3.39E-26 |
| KRAS | -2.989 | 3.37E-24 |
| PDGF BB | -3.101 | 4.29E-36 |
| TNF | -3.435 | 2.34E-29 |
| HRAS | -4.235 | 8.86E-37 |

Genes with adjusted p-value<0.05

**b**

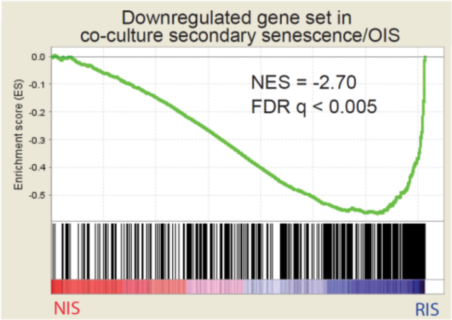


Leading edge genes: STC1, MMP1, SERPINB2, CDCP1, NRG1, SLC16A6, TFPI1, TMEM158, CSF3, IL8, MT2A, HMGA1, IL1B

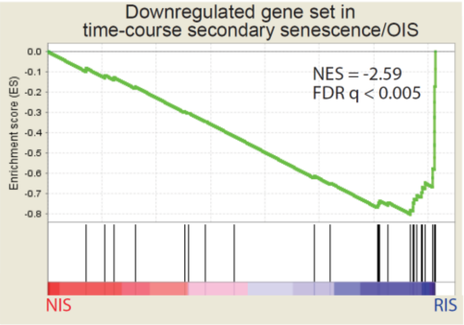


Leading edge genes: MMP3, STC1, MMP1, SERPINB2, SLC16A6, TFP12, IL2B,

**c**

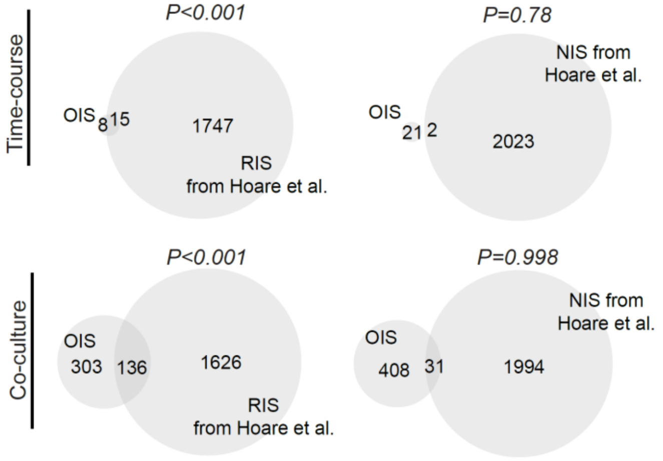

**Supplementary Fig. 2. Primary senescence comprises RIS signature. a.** IPA analysis of the two senescence clusters from time-course and co-culture scRNA-seq. Red indicates activated upstream regulator and blue indicates inhibited upstream regulator. **b.** GSEA was used to assess the enrichment of secondary and primary senescence (OIS) DE genes in Hoare et al.'s NIS and RIS log2FC preranked genes. NES and FDR are shown. **c.** Venn diagrams overlapping expression signatures from top panel: time-course and bottom panel: co-culture experiments with NIS signature genes (OIS: OIS/Secondary senescence upregulated genes; NIS: Hoare et al.'s NIS/RIS upregulated genes; RIS: Hoare et al.'s RIS/NIS upregulated genes)

Fig. S3

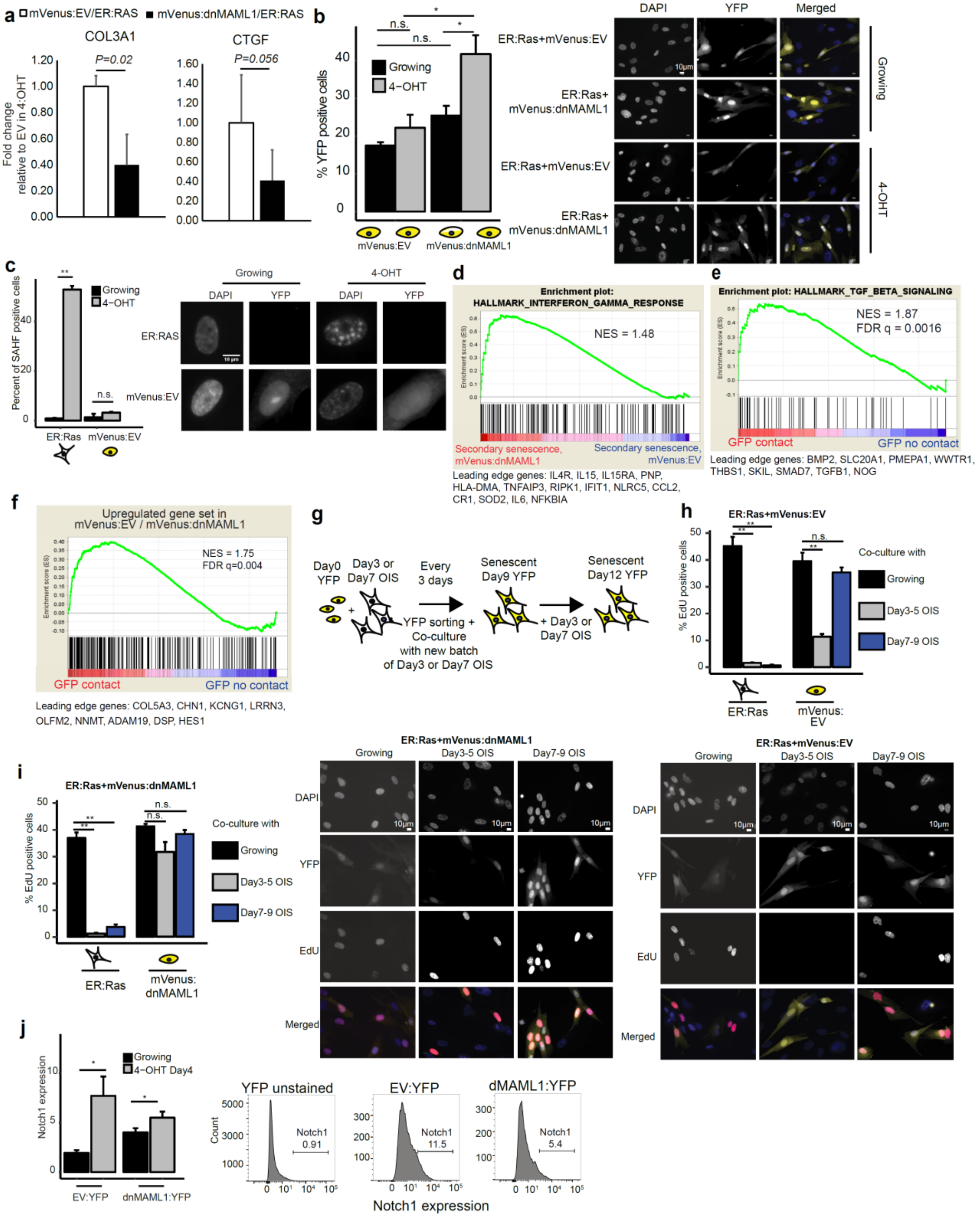
a
mVenus:EV/ER:RAS
mVenus:dnMAML1/ER:RAS
COL3A1
P=0.02
CTGF
P=0.056
Fold change relative to EV in 4-OHT
b
% YFP positive cells
Growing
4-OHT
n.s.
mVenus:EV
mVenus:dnMAML1
DAPI
YFP
Merged
ER:Ras+mVenus:EV
ER:Ras+ mVenus:dnMAML1
Growing
4-OHT
c
Percent of SAHF positive cells
ER:Ras
mVenus:EV
d
Enrichment plot: HALLMARK_INTERFERON_GAMMA_RESPONSE
NES = 1.48
Secondary senescence, mVenus:dnMAML1
Secondary senescence, mVenus:EV
Leading edge genes: IL4R, IL15, IL15RA, PNP, HLA-DMA, TNFAIP3, RIPK1, IFIT1, NLRC5, CCL2, CR1, SOD2, IL6, NFKBIA
e
Enrichment plot: HALLMARK_TGF_BETA_SIGNALING
NES = 1.87
FDR q = 0.0016
GFP contact
GFP no contact
Leading edge genes: BMP2, SLC20A1, PMEPA1, WWTR1, THBS1, SKIL, SMAD7, TGFB1, NOG
f
Upregulated gene set in mVenus:EV / mVenus:dnMAML1
NES = 1.75
FDR q=0.004
GFP contact
GFP no contact
Leading edge genes: COL5A3, CHN1, KCNG1, LRRN3, OLFM2, NNMT, ADAM19, DSP, HES1
g
Day0 YFP
Day3 or Day7 OIS
Every 3 days
YFP sorting + Co-culture with new batch of Day3 or Day7 OIS
Senescent Day9 YFP
+ Day3 or Day7 OIS
Senescent Day12 YFP
h
ER:Ras+mVenus:EV
% EdU positive cells
Co-culture with
Growing
Day3-5 OIS
Day7-9 OIS
ER:Ras
mVenus: EV
i
ER:Ras+mVenus:dnMAML1
mVenus: dnMAML1
ER:Ras+mVenus:EV
EdU
j
Notch1 expression
Growing
4-OHT Day4
EV:YFP
dnMAML1:YFP
YFP unstained
EV:YFP
dMAML1:YFP
Count
Notch1
0.91
11.5
5.4

**Supplementary Fig. 3. SAHF formation in primary and secondary senescence. a.** Bar plot showing the expression of *CTGF* (n=6) and *Col3A1* (n=3) genes in EV or dnMAML1 cells compared to ER:Ras senescent cells by qPCR. (COL3A1: t=5.3405, df=2.4861, p=0.02; CTGF: t=2.2104, df=8.4894, p=0.056 using unpaired Student's t- test). Error bars represent SEM. **b.** Bar plot denoting the proportion of growing (black) or senescent (grey) mVenus cells with dnMAML1 or EV as proportion of all cells scored. Error bars are displayed as SEM; $F[3,8] = 10.05$, $p<0.05$ using one-way ANOVA with Tukey's test (n=3 for each condition). Images of mVenus cells and cells stained with DAPI were shown on the right of the barplot. Scale bar 10µm. **c.** Primary senescence induced by Ras showed significant SAHF formation but not in secondary senescent cells as identified by unpaired Student's t-test. (ER:Ras t=-34.05, df=2.12, ** p<0.01; mVenus:EV t=-1.23, df=2.28, p=0.32; n=3 for each condition). Representative images are shown on the right. **d.** GSEA pre-ranked test showed the enrichment of interferon gamma response in mVenus:dnMAML1 identified as secondary senescence using scmap. **e.** GSEA pre-ranked test showed the enrichment of TGF-beta signaling in GFP contact cells in comparison to GFP no contact cells. mVenus:dnMAML1 identified as secondary senescence using scmap. **f.** GSEA pre-ranked test showed the enrichment of mVenus:EV signature genes in GFP contact/GFP no contact upregulated gene set. **g.** Schematic representation of co-culturing mVenus cells with Day3 or Day7 OIS cells. **h.** Bar plot showing EdU incorporation in OIS or mVenus:EV cells in growing (black), co- culture with Day3 OIS (grey) or Day7 OIS cells (blue). Error bars are displayed as SEM; $F[5,18] = 144.4$, $p<0.001$ using one-way ANOVA with Tukey's test (n=3 for each except for Day3 OIS (n=6)). Representative images are shown at the bottom of the bar plot. Scale bar 10µm. **i.** Bar plot showing EdU incorporation in OIS or mVenus:dnMAML1 cells in growing (black), co-culture with Day3 OIS (grey) or Day7 OIS cells (blue). Error bars are displayed as SEM. $F[5,24] = 58$, $p<0.001$ using one-way ANOVA with Tukey's test (n=3 for all conditions except for Day3 OIS (n=6)). **p<0.001, *p<0.05. Representative images are shown on the right of the bar plot. Scale bar 10µm. **j.** Barplot showing the significant upregulation of Notch1 on mVenus:EV and mVenus:dnMAML1 cells compared to growing (mVenus:EV t = -3.27, df = 2.01, p-value = 0.041; mVenus:dnMAML1 t = -3.29, df =3.03, p-value = 0.023 using one-sided t- test). Error bars represent SEM. Representative FACs plots showing Notch1 staining of YFP uninduced fibroblasts and YFP:EV and YFP:dnMaml1 at 4 days of co-culture.

### Supplementary Fig. 4

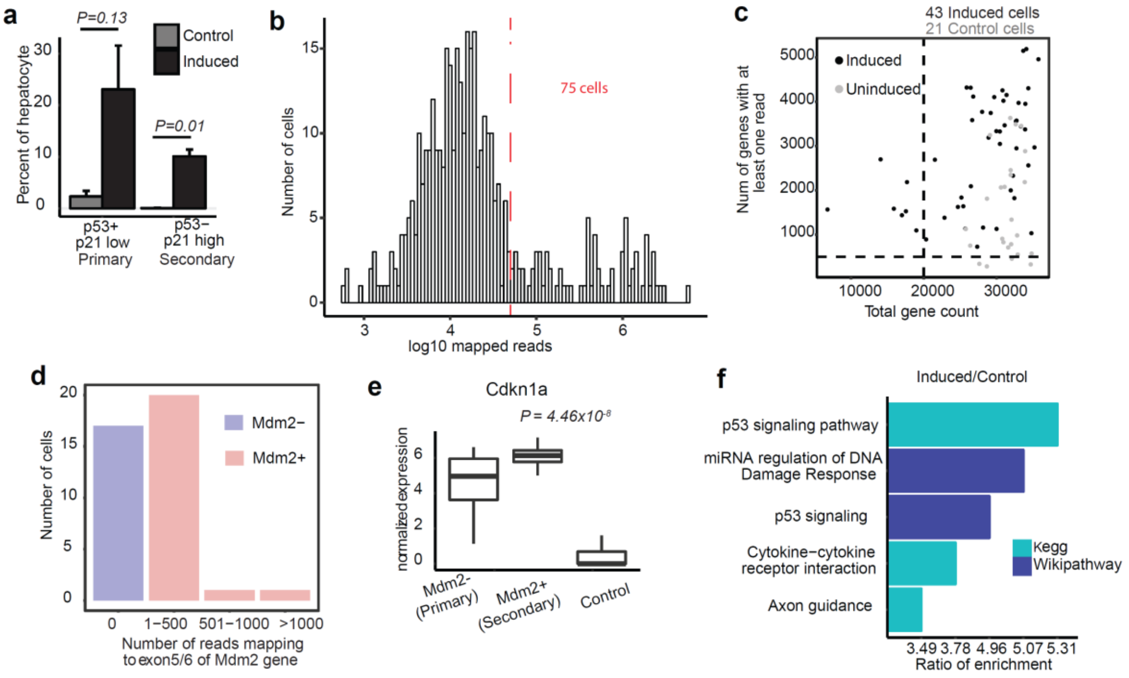

**Supplementary Fig. 4. Quality control and processing of single cells. a.** Percent of primary and secondary hepatocytes (Primary: t=2.4241, df=2.0641, p-value = 0.1324; Secondary: t=7.7563, df=2.0053, p=0.0161 using unpaired Student's t-test). **b**. Histogram with the number of induced and control cells is plotted against log mapped reads. 75 single cells with at least 50,000 aligned reads are downsampled to 50,000 reads. **c.** Dot plot with the number of genes with at least one read to total gene count for induced (black) and control (grey) cells. Cells with a total gene count of more than 20,000 and 500 genes detected were retained. **d.** 17 Mdm2- cells were identified as cells with no reads mapping to exon5/6 of *Mdm2* gene and 22 Mdm2+ cells contained reads mapping to the exons. **e.** Box plots showing the expression of Cdkn1a in induced cells relative to control ($p=4.46x10^{-18}$). The top and bottom bounds of the boxplot correspond to the 75 and 25th percentile, respectively. **f**. p53 signaling and cytokine-cytokine receptor interaction were enriched in induced cells versus control, confirming the senescence phenotype.

**Table S1.**
Type or paste caption here. Create a page break and paste in the Table above the caption.

<insert Table S1 here followed by a page break >

**Table S2.**
Type or paste caption here. Create a page break and paste in the Table above the caption.

<insert Table S2 here followed by a page break >
